# Pairing extinction training with vagus nerve stimulation (VNS) reduces drug-seeking by altering activity in afferents to the medial prefrontal cortex

**DOI:** 10.64898/2025.12.03.691662

**Authors:** Christopher Driskill, Lily Vu, Sofia Jalilvand, Frank Salazar, Laney Waydick, Neha Suji, Sanjana Tata, Aamna Khan, Zarin Hasan, Ria Nuna, Zara Kanwal, Neissa Molin, Sven Kroener

**Affiliations:** University of Texas at Dallas, Department of Neuroscience, Richardson, TX, USA

**Keywords:** extinction, parvalbumin, amygdala, hippocampus, cocaine

## Abstract

Relapse triggered by drug-associated cues or stress remains a major challenge in treating substance use disorders (SUDs), as re-exposure reliably provokes craving and reinstatement of drug seeking. Extinction-based interventions can reduce cue reactivity, yet extinction learning is often weak or context-dependent, limiting its clinical impact. Vagus nerve stimulation (VNS) enhances learning-related plasticity via widespread engagement of neuromodulatory systems and cortical circuits. Recent preclinical work shows that pairing extinction with VNS facilitates extinction learning and reduces cue-induced reinstatement of cocaine seeking, suggesting translational potential as an adjunct to exposure-based therapies. However, the circuit-level mechanisms underlying these effects remain unclear. To address this gap, we examined how VNS paired with extinction alters activity in medial prefrontal cortex (mPFC) networks that regulate drug seeking, focusing specifically on afferent projections from the basolateral amygdala (BLA) and ventral hippocampus (vHPC). Male rats received retrograde AAV-eGFP infusions into either the prelimbic (PL) or infralimbic (IL) cortex to label upstream projections, followed by cocaine self-administration, extinction training with VNS or sham stimulation, and cue-induced reinstatement. cFos immunolabeling in the BLA and vHPC revealed pathway-specific modulation: VNS decreased overall BLA activity and reduced activation of BLA to IL projections, but increased activation of BLA to PL projections. In the vHPC, VNS selectively decreased activation of vHPC to IL projections without affecting PL-projecting neurons. Because these pathways synapse onto parvalbumin interneurons (PVIs) in the mPFC, we quantified PVI activation and found that VNS decreased overall prefrontal cFos expression, but increased PVI activity in the PL, and decreased PVI activity in the IL. Together, these results demonstrate that VNS paired with extinction reshapes prefrontal–amygdala–hippocampal circuits in a pathway-specific manner, potentially modulating feed-forward inhibition to reduce relapse-like behavior. These findings support VNS as a promising strategy to strengthen extinction learning and improve treatment outcomes in SUD.

## Introduction

In patients with substance use disorder (SUD) re-exposure to drug-associated cues or stress induces craving and relapse (O’Brien, Childress et al. 1998, Sinha, Fuse et al. 2000). In extinction training the reinforcing consequences of drug taking are removed to reduce cue-reactivity and drug-seeking (Conklin and Tiffany 2002, Havermans and Jansen 2003, Millan, Marchant et al. 2011, Phillips, Epstein et al. 2014, Perry and Lawrence 2017). Facilitating extinction processes in order to prevent relapse is therefore clinically important (Taylor, Olausson et al. 2009). Vagus nerve stimulation (VNS) is FDA-approved for use in epilepsy and depression (Rush, Marangell et al. 2005, Nemeroff, Mayberg et al. 2006), and it might be repurposed as an adjuvant in exposure-based therapies for the treatment of SUDs. VNS causes the release of several neuromodulators which modulate cortical plasticity (Krahl and Clark 2012, Cao, Lu et al. 2017), and this plasticity can be used to increase learning and memory in both rats (Clark, Smith et al. 1998, Cao, Wang et al. 2016, Driskill, Childs et al. 2022) and humans (Clark, Naritoku et al. 1999, Sun, Perakyla et al. 2017). VNS-induced plasticity facilitates extinction and reduces cue-induced reinstatement of drug-seeking in cocaine self-administering rats (Childs, DeLeon et al. 2017, Childs, Kim et al. 2019, Driskill, Childs et al. 2024). These changes correlate with altered cellular activity in a network that involves the medial prefrontal cortex (mPFC), the basolateral amygdala (BLA), and the Nucleus accumbens (NAc) (Childs, DeLeon et al. 2017).

In rodents the mPFC receives inputs from the amygdala and hippocampus, as well as other limbic structures, positioning it to integrate information regarding salience, value, and contextual cues associated with both appetitive and aversive outcomes (Peters, Kalivas et al. 2009, Gourley and Taylor 2016). Within the mPFC, the prelimbic (PL) and infralimbic (IL) subregions are believed to serve largely opposing roles in regulating conditioned responses to both rewarding and aversive stimuli: The PL is implicated in the formation and expression of conditioned responses, and in the context of drug-seeking it drives reinstatement via its projection to the Nucleus accumbens core (NAc core). Conversely, the IL is important for the extinction of conditioned responses, inhibiting cocaine-seeking through its projections to the Nucleus accumbens shell (NAc shell) and/or to the PL (Peters, Kalivas et al. 2009, Peters, Pattij et al. 2013, Muller Ewald, De Corte et al. 2019). Here we further explored the network mechanisms through which VNS reduces drug-seeking behavior. We focused on VNS-induced changes in afferents to the mPFC from the BLA and the ventral hippocampus (vHPC), two areas that interact with the mPFC to regulate appetitive associative learning, memory retrieval, and extinction. Our goal was to determine whether pairing extinction training with VNS would differentially activate projections from these upstream regions to the PL and IL during reinstatement. To test this idea, we combined retrograde tracing from the IL and PL with immunohistochemical detection of cFos (Cruz, Javier Rubio et al. 2015) following cue-induced reinstatement in male rats that had received VNS or Sham-stimulation during extinction. In addition, in the mPFC we analyzed VNS-induced changes in cFos expression in parvalbumin-expressing interneurons (PVIs), which are targets of projections from the BLA and vHPC. Our results reveal a complex pattern of changes, with both global changes in cFos activity, as well as pathway-specific increases and decreases in projections to the IL and PL. In the mPFC, we found a decrease in cFos colabeling in PVI in the IL, as well as an opposite increase of cFos colabeling in PVI in the PL, indicating differential modulation of inhibition of mPFC networks by VNS. These results shed light on pathway-specific changes in the activation of areas that project to the mPFC that can be affected by pairing extinction with VNS.

## Methods

### Subjects

Male Sprague-Dawley rats (Taconic, Germantown, NY) were individually housed and kept on a 12-hr. reverse light/dark cycle, with free access to food and water until surgery, when food was restricted to 25 g/day standard rat chow. At the time of surgery rats were at least 90 days old (250-300g). All protocols were approved by the IACUC of The University of Texas at Dallas and were conducted in compliance with the NIH Guide for the Care and Use of Laboratory Animals.

### Retrograde tracing

To visualize mPFC projections we infused a retrograde AAV expressing eGFP (*pENN.AAV.hSyn.HI.eGFP-Cre.WPRE.SV40*; 1×10^13^ vg/mL; Addgene viral prep #105540-AAVrg) bilaterally (0.4 μl each hemisphere, rate of 80nL/min) into either the PL (+3.0 A/P, ±0.6 M/L, -3.9 D/V) or the IL (+3.0 A/P, ±0.6 M/L, -5.4 D/V) (Paxinos 2007). Rats recovered for 7 days prior to VNS cuff and catheter implantation surgery.

### Drug Self-Administration and Extinction Training

Drug self-administration and extinction training were performed as previously described (Childs, DeLeon et al. 2017, Driskill, Childs et al. 2024). Rats were anesthetized and implanted with a catheter in the right external jugular vein for drug administration. During the same surgery, a custom-made cuff electrode was placed around the left vagus nerve for the delivery of VNS (Childs, Alvarez-Dieppa et al. 2015). Seven days following surgery, rats were trained in a single overnight session to self-administer food pellets (45 mg, Bio Serv, Flemmington, NJ) in an operant conditioning chamber (Med Associates, Saint Albans, VT). Drug self-administration training took place in the same chamber, which was equipped with two levers, a house light, a cue light, and a tone. Each active lever press produced a 0.05 ml infusion of 2.0 mg/mL cocaine (NIDA Drug Program) in saline, and the presentation of drug-paired cues (illumination of the light over the active lever and the presentation of a 2900 Hz tone), followed by a 20 s timeout. Self-administration sessions ended after 2 hours. Both right and left levers were available for the duration of the session and drug-seeking behavior was quantified as active lever presses. Rats self-administered cocaine for 15-18 days, with a minimum criterion of at least 20 infusions per session. Subjects then underwent 10 days of extinction training in which lever presses on the previously active lever no longer produced cocaine or presentation of drug-paired cues. During extinction training rats received either sham-stimulation or non-contingent VNS (0.4 mA, 500 µs pulse width at 30 Hz, stimulation cycle of 30 sec on, every 5 min) for the duration of the training session. After 10 days of extinction training, drug-seeking behavior was reinstated by presentation of the drug-associated cues in the operant conditioning chambers. During the reinstatement session presses on the previously active lever led to presentation of the drug-associated tone and light but did not result in drug delivery or VNS.

### Immunohistochemistry

We used immunohistochemistry to colocalize AAVrg-eGFP labeled cells with the activity marker cFos in VNS- and Sham-stimulated rats following cue-induced reinstatement. Sixty minutes after the reinstatement session, rats were anesthetized with an overdose of urethane (3 g/kg i.p.) and transcardially perfused with room temp 1x PBS followed by 4% paraformaldehyde in 1x PBS (4°C, pH 7.4). Brains were postfixed in PFA with 30% sucrose for 3 hrs. and were then transferred to 30% sucrose in PBS for approximately 18 hrs. at 4°C. Coronal slices (40 μm) were cut on a freezing microtome and collected in PBS containing 0.01% NaN3 as a preservative. To determine VNS-induced changes in cFos expression and the colocalization of retrogradely labeled GFP+ cells with cFos we analyzed the ventral subiculum and CA1 area of the ventral hippocampus in slices between -4.68 and -5.40 relative to bregma. Similarly, we analyzed cFos expression and GFP+/cFos colocalization in nuclei in the Basolateral Amygdala Complex, including the ventrolateral part of lateral nucleus (LaVL), the ventromedial part of lateral nucleus (LaVM), the anterior part of basolateral nucleus (BLa), and the posterior part of basolateral nucleus (BLp) in slices between bregma -2.52 and -3.24. These nuclei are referred here collectively as “BLA” and the results of their analyses were pooled together.

Free-floating sections containing BLA and vHPC were incubated in guinea pig monoclonal recombinant anti-cFos (Synaptic Systems, Cat# 226 308; RRID:AB_2905595; 1:10,000 working dilution) in PBS with 0.5% Triton and 2% normal goat serum (Thermo Fisher Scientific,) for 36 hrs. at 4°C. Sections were washed at least 3 times for 10 min each in PBS before they were incubated in donkey monoclonal anti-Guinea pig Alexa 647 (Thermo Fisher Scientific, Cat# A-21450, RRID:AB_2535867, 1:5000 working dilution) for 2 hrs. at room temperature in 0.5% Triton-X and 2% normal goat serum in PBS. Sections were washed 3 times in PBS before they were mounted and cover-slipped using Prolong Gold Antifade with DAPI (Invitrogen, Grand Island, NY). For each animal a minimum of 4 sections of the mPFC, vHPC and BLA were imaged as z-stacks (3-micron step size) on a confocal microscope (FV3000, Olympus Corporation, Tokyo, Japan) with a 10x objective. Images were converted to Imaris file format and analyzed by an experimenter blind to the treatment conditions.

To assess changes in overall cFos expression and cFos expression within parvalbumin-expressing interneurons (PVIs), we analyzed coronal sections of the mPFC (IL and PL) spanning Bregma +3.72 to +2.52. cFos immunohistochemistry was performed as described above. For colabeling of cFos and parvalbumin, sections were incubated with a rabbit anti-PV primary antibody (Swant, Cat# PV27, RRID:AB_2631173; 1:2000 working dilution) together with the anti-cFos antibody during the primary incubation. Following washes, sections were incubated with a goat anti-rabbit Alexa Fluor 546 secondary antibody (Thermo Fisher Scientific, Cat# A-11010, RRID:AB_2534077; 1:1000 working dilution).

### Data analysis

All statistical analyses were performed in GraphPad Prism 7.0.5 (GraphPad Software). We compared lever presses on the first day of extinction and during reinstatement with a one-way ANOVA, Post hoc analyses of main effects used Tukey’s multiple-comparison tests. Unpaired *t*-tests were used to compare differences in GFP+ neurons, cFos+ neurons, and cFos expression in GFP+ neurons. To compare changes in the mPFC in cFos+ neurons and cFos expression in PVIs we used separate two-way ANOVAs with factors of treatment and region. Post hoc analysis was performed with an uncorrected Fisher’s Least Significant Difference (LSD) test.

## Results

### Vagus nerve stimulation during extinction reduces cue-induced reinstatement

Separate groups of rats received infusions of retrograde AAV expressing eGFP into either the infralimbic cortex (IL) or prelimbic cortex (PL). These rats were then trained to self-administer cocaine for 15-18 days, followed by 10 days of extinction training paired with either vagus nerve stimulation (VNS) or Sham stimulation. Twenty-four hours after the last day of extinction training, drug-seeking behavior was reinstated in a cued reinstatement session by presenting the conditioned drug cues (Figure 1). Figure 1A shows lever presses during the last 10 days of drug-self administration, during the 10 days of extinction, and the reinstatement session in rats that received retro-eGFP infusions into the IL. We compared behavior between rats that received Sham stimulation (n=8) or VNS (n=8) during the extinction period. A one-way ANOVA found a significant effect of treatment on lever presses during the first day of extinction (F_(3, 28)_ = 14.24, p < 0.0001, Figure 1B). Post hoc analysis with Tukey’s multiple comparisons test showed a significant decrease in active lever presses (p = 0.0009), but no difference in the number of inactive lever presses (p = 0.7856). Similarly, a one-way ANOVA for responses during the cue-induced reinstatement session found a significant effect of treatment on lever presses (F_(3,28)_ = 9.087, p = 0.0002, Figure 1C). Post hoc analysis with Tukey’s multiple comparisons test showed a significant decrease in active lever presses in animals that received VNS (p = 0.0212), but no difference in the number of inactive lever presses (p = 0.9923). We performed the same analyses for drug-seeking behavior during the extinction period and the reinstatement session, respectively, in rats that received an AAV infusion of retro-eGFP into the PL (Figure 1D) comparing lever presses between rats that received VNS (n=7) or Sham-stimulation (n=7). A one-way ANOVA found a significant effect of treatment on lever presses during the first day of extinction (F_(3, 24)_ = 19.58, p < 0.0001, Figure 1E). Post hoc analysis with Tukey’s multiple comparisons test showed a significant decrease in active lever presses in animals that received VNS (p < 0.0001), but no difference in the number of inactive lever presses (p = 0.5350). A one-way ANOVA for lever presses during the cue-induced reinstatement session found a significant effect of treatment (F_(3,24)_ = 7.925, p = 0.0008, Figure 1F). Post hoc analysis with Tukey’s multiple comparisons test showed a significant decrease in active lever presses in animals that received VNS (p = 0.0143), but no difference in the number of inactive lever presses (p = 0.9956).

**Figure 1.**
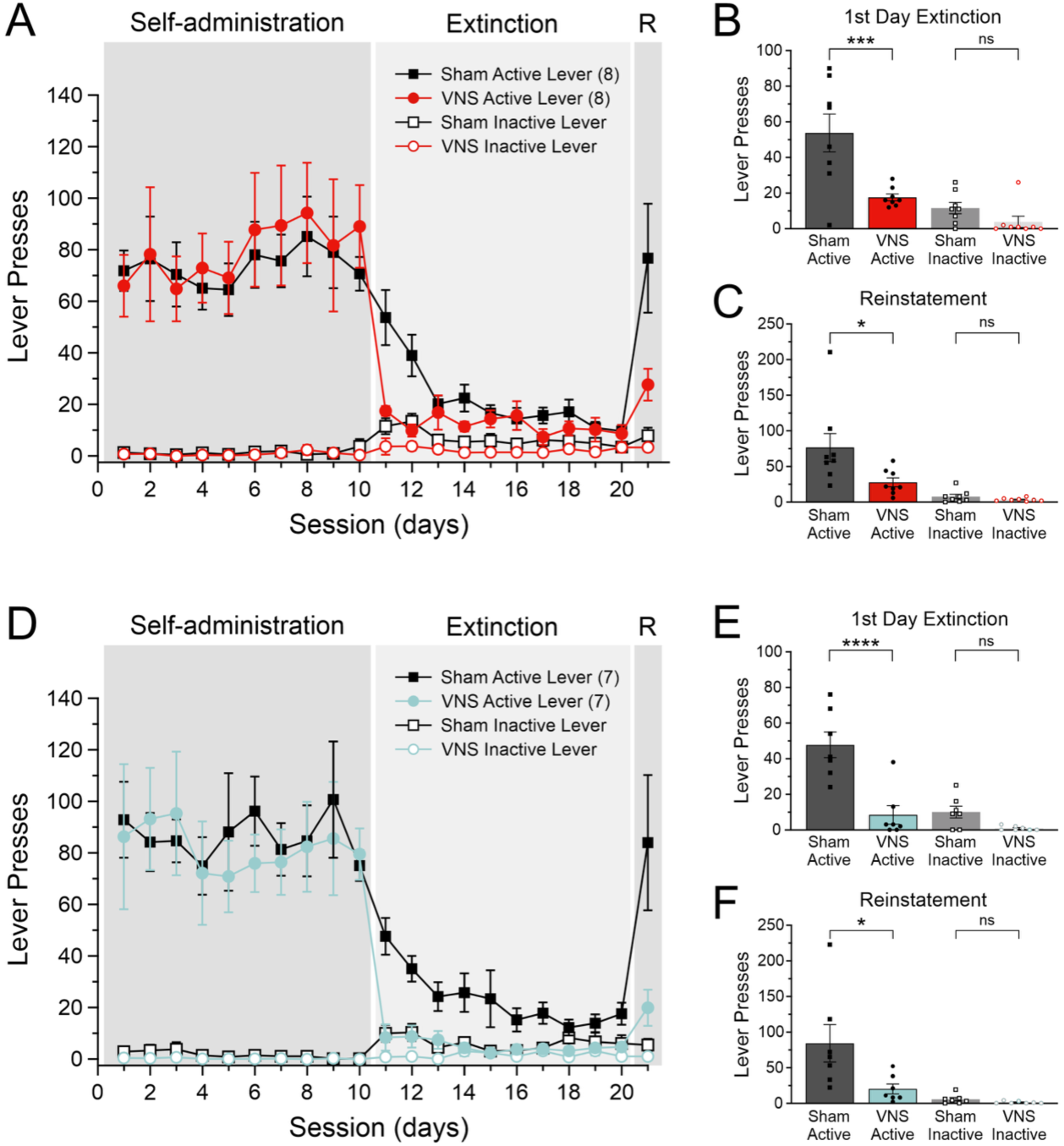
Vagus nerve stimulation (VNS) facilitates extinction from cocaine seeking and reduces cue-induced reinstatement. A) Active (solid symbols) and inactive lever presses (open symbols) in rats that received infusions of a retrograde AAV into the infralimbic cortex (IL) during cocaine self-administration, extinction, and cue-induced reinstatement (R). Rats received either VNS (red symbols, n=8) or sham stimulation (black symbols, n=8) during extinction on days 11–20. B) VNS-treated rats displayed reduced active lever presses during the first day of extinction. C) Responses on the previously active lever during cue-induced reinstatement are significantly reduced in VNS-treated rats. D) Active (solid symbols) and inactive lever presses (open symbols) in rats that received infusions of the retrograde AAV into the prelimbic cortex (PL) during cocaine self-administration, extinction, and cue-induced reinstatement (R). Rats in this cohort also received either VNS (Teal symbols, n=7) or sham-stimulation (Black symbols, n=7) during extinction training. E) VNS-treated rats with PL infusions also showed accelerated extinction on the first day of extinction, F) and reduced responding at the active lever during cue-induced reinstatement. P values are (*) <0.05, (***) <0.001, and (****) <0.0001.

### Distribution of neurons projecting to the mPFC

Figure 2 shows the brain-wide distribution of retrogradely eGFP-labeled cells in major nuclei of the rat brain that project to either the IL (red symbols) or the PL (teal symbols). For each infusion group (PL or IL) we took sections from four rats (2 VNS- and 2 Sham-treated rats each) to map out the distribution of eGFP-expressing neurons. For each brain region we took the average number of eGFP-expressing cells across the four brains and placed one dot for approximately every 5-10 cells. Significant numbers of cells were found in midline thalamic nuclei, the BLA, the vHPC, and the major sources of neuromodulatory inputs to the mPFC, including the ventral tegmental area (VTA), midline raphe nuclei, and the locus coeruleus (LC). In this report we focus on VNS modulation of the projections from the CA1 region of the vHPC, and the BLA.

**Figure 2:**
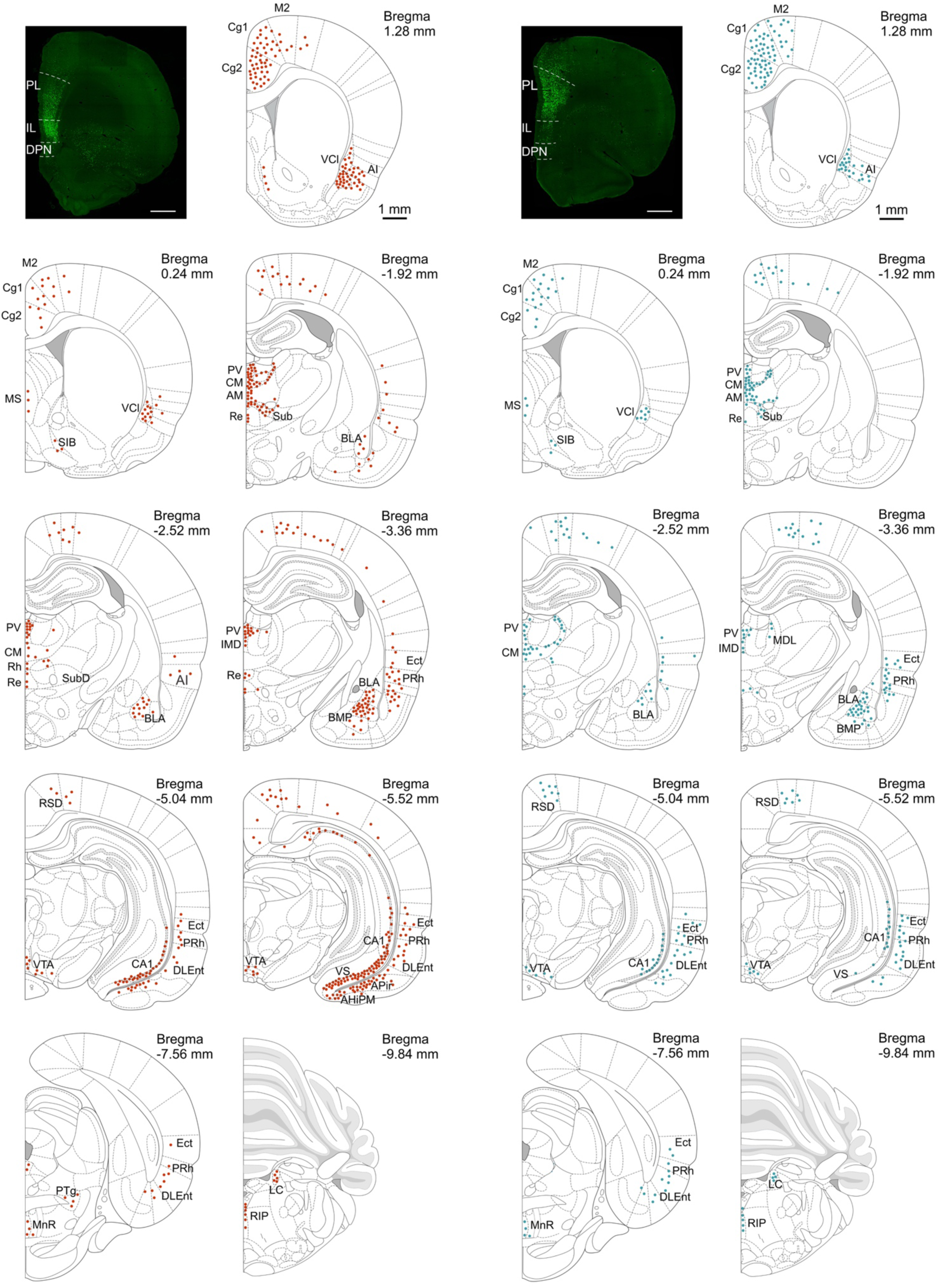
Photomicrographs of infusions sites of a retrograde AAV expressing eGFP in the infralimbic cortex (IL, left) and the prelimbic cortex (PL, right) and corresponding distribution of retrogradely labeled cells in major brain nuclei (IL, red symbols; PL teal symbols). Each dot represents about 5-10 cells; averages for 4 rats (2 Sham and 2 VNS) for each infusion site. Abbreviations: AHiPM, Amygdalohippocampal area, posteromedial part; Ai, Agranular insular cortex; AM, Anteromedial thalamic nucleus; APir, Amygdalopiriform transition area; BLA, Basolateral amygdala complex; BMP, Basomedial amygdaloid nucleus, posterior part; CA1, Cornu Ammonis area 1 of the hippocampus; Cg1, Cingulate cortex, area 1; Cg2, Cingulate cortex, area 2; CM, Central medial thalamic nucleus; DLEnt, Dorsolateral entorhinal cortex; Ect, Ectorhinal cortex; IMD, Intermediodorsal thalamic nucleus; LC, Locus coeruleus; M2, Secondary motor cortex; MDL, Mediodorsal thalamic nucleus, lateral part; MnR, Median raphe nucleus; MS, Medial septum; PRh, Perirhinal cortex; PTg, Pedunculotegmental nucleus; PV, Paraventricular thalamic nucleus; Re, Reuniens thalamic nucleus; RIP, Raphe interpositus neucleus; RSD, Retrosplenial dysgranular cortex; SiB, Substantia innominata, basal part; Sub, Submedius thalamic nucleus; VCl, ventral part of claustrum; VS, Ventral subiculum; VTA, Ventral tegmental area.

### Effect of VNS on projections from the BLA to the mPFC

We analyzed cFos expression during drug-seeking and colocalization of cFos in IL-projecting neurons in the basolateral complex in brains from the same rats receiving VNS (n=8) or Sham-stimulation (n=8) (Figure 3A-E). Separate unpaired t-tests showed no difference in the number of GFP+ cells (t_(14)_ = 0.3803, p = 0.7094, Figure 3C); however, there was a significant overall decrease in cFos+ cells (t_(14)_ = 2.836, p = 0.0132, Figure 3D), and a decrease in the percentage of GFP+ cells that expressed cFos (t_(14)_ = 4.885, p = 0.0002, Figure 3E). We then performed the same comparisons in VNS- and Sham stimulated rats that received retro-eGFP infusions into the PL (Figure 3F-J). Separate unpaired t-tests found no difference in the number of GFP+ cells (t_(12)_ = 0.6664, p = 0.5177, Figure 3H), but there was an overall decrease in cFos+ cells in VNS-treated rats (t_(12)_ = 2.653, p = 0.0211, Figure 3I), and an increase in the percentage of GFP+ cells that expressed cFos in VNS-treated rats (t_(12)_ = 2.552, p = 0.0254, Figure 3J). Taken together, these data suggest that pairing extinction with VNS selectively modulates neuronal activity in two pathways from the BLA to the IL and PL, respectively, in an opposite manner.

**Figure 3:**
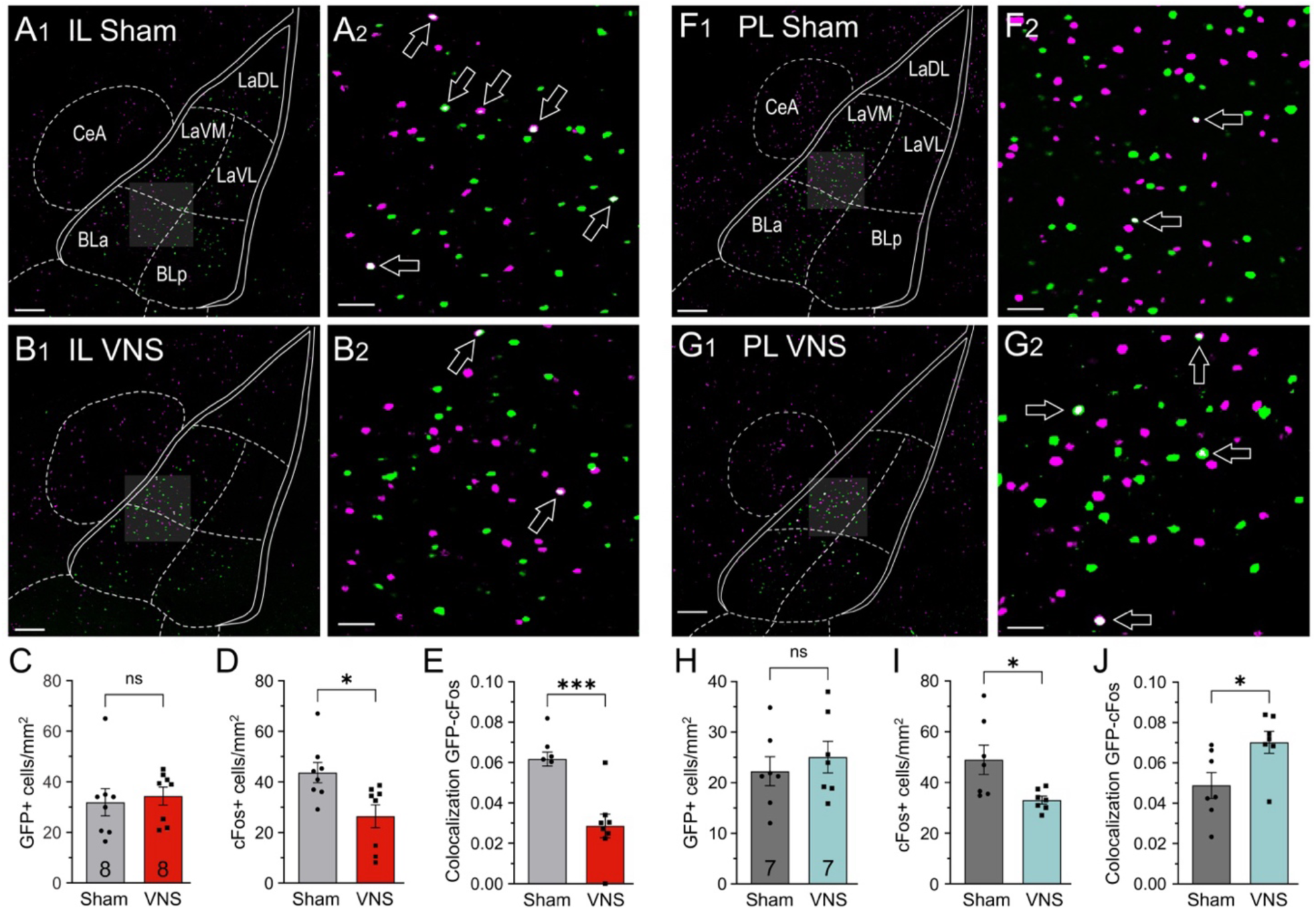
VNS differentially modulates cFos expression following reinstatement in IL- and PL-projecting neurons of the basolateral amygdala (BLA). A, B) GFP-positive cells (green) following infusion of a retro-AAV into the IL and cFos (pink) in the BLA of rats that received either Sham-stimulation (n=8), (Figure 3A) or VNS (n=8) during extinction (Figure 3B). A_2_, B_2_). Arrows indicate GFP+ cells colocalized with cFos (white). C) VNS and Sham-stimulated rats did not differ in the number of GFP+ cells in the BLA. D) VNS reduced overall cFos expression in the BLA. E) VNS also reduced cFos expression in IL-projecting (GFP+) neurons. F, G) GFP-positive cells following infusion of the retro-AAV into the PL and cFos in the BLA of rats receiving either Sham-stimulation (n=7) (Figure 3F), or VNS (n=7) during extinction (Figure 3G). H) VNS and Sham-stimulated rats did not differ in the number of GFP+ cells in the BLA. I) VNS also reduced overall cFos expression in the BLA in this cohort. J) In contrast to its effect on IL-projecting neurons, VNS selectively increased cFos labeling in PL-projecting neurons. Scale bars represent 200 µm in A_1_, B_1_, F_1_ and G_1_, and 100 µm in A_2_, B_2_, F_2_ and G_2_. *P* values are (*) <0.05, and (***) <0.001.

### Effect of VNS on ventral hippocampal projections to the mPFC

In rats that received AAV infusions of retro-eGFP into the IL we measured the number of cFos+ cells in the ventral subiculum and CA1 region of the vHPC as an indicator of cellular activity during cue-induced reinstatement in Sham-stimulated rats (n=8) and rats given VNS (n=8) (Figure 4A-E). Similarly, we measured the colocalization of IL-projecting (GFP+) neurons with cFos as a measure of activation of this pathway. Separate unpaired t-tests found no difference in the number of GFP+ cells (t_(14)_ = 1.574, p = 0.1377, Figure 4C), no difference in the total number of cFos+ cells (t_(14)_ = 0.01091, p = 0.9914, Figure 4D), and a decrease in the percentage of GFP+ cells that expressed cFos (t_(14)_ = 2.303, p = 0.0371, Figure 4E) between Sham- and VNS-treated rats.

**Figure 4:**
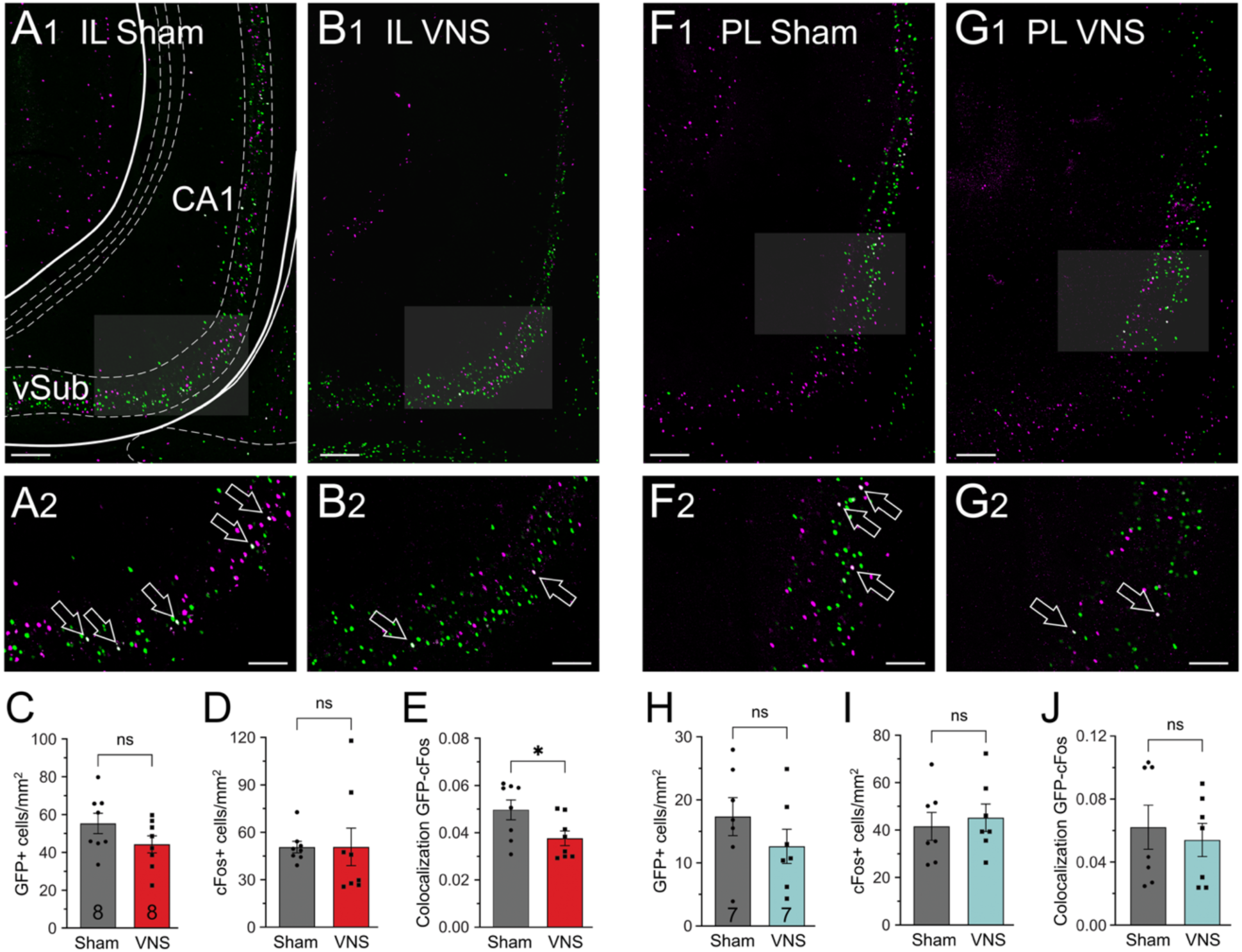
VNS modulates cFos expression following reinstatement differentially in IL- and PL-projecting neurons of the ventral hippocampus (vHPC). A, B) GFP-positive cells (green) following retro-AAV infusion into the IL and cFos (pink) in the vHPC of rats receiving either Sham-stimulation (n=8), (Figure 4A) or VNS (n=8) during extinction, (Figure 4B). A_2_, B_2_) Arrows indicate GFP+ cells colocalized with cFos (white). C) VNS and Sham-stimulated rats did not differ in the number of GFP+ cells in the vHPC. D) VNS did not significantly affect overall cFos expression in the BLA. E) VNS also reduced cFos expression in IL-projecting (GFP+) neurons. F, G) GFP-positive cells following infusion of the retro-AAV into the PL and cFos in the vHPC of rats receiving either Sham-stimulation (n=7), F) or VNS (n=7) during extinction. H) VNS and Sham-stimulated rats did not differ in the number of GFP+ cells in the vHPC. I) VNS did not affect overall vHPC cFos expression. J) VNS did not alter cFos labeling in PL-projecting neurons. Scale bars represent 200 µm in A_1_, B_1_, F_1_ and G_1_, and 100 µm in A_2_, B_2_, F_2_ and G_2_. *P* values are (*) <0.05.

Next, we performed the same analyses in rats that received AAV infusions of retro-eGFP into the PL. We compared eGFP and cFos expression, as well as their colocalization between rats that received Sham stimulation (n=7) or VNS (n=7) (Figure 4F-J). Separate unpaired t-tests found no difference in the number of GFP+ cells (t_(12)_ = 1.166, p = 0.2661, Figure 4H) and no difference in cFos+ cells (t_(12)_ = 0.4423, p = 0.6661, Figure 4I); and there was no difference in the percentage of GFP+ cells that expressed cFos in VNS treated rats (t_(12)_ = 0.4659, p = 0.6497, Figure 4J). These results indicate pathway-specific modulation of the projection from the vHPC to the IL by extinction paired with VNS.

### VNS modulation of mPFC PVI activity during drug-seeking

Projections from the vHPC and BLA to the mPFC make connections with PVIs that are powerful regulators of networks and behavioral output (McGarry and Carter 2016, Marek, Jin et al. 2018, Aleman-Andrade, Witter et al. 2025). We stained slices of the mPFC for PV and cFos to determine activity of PVIs during cue-induced reinstatement in rats that received VNS (n=15) or Sham-stimulation (n=14) (Figure 5). We first quantified total cFos expression comparing PL to IL and Sham-stimulation to VNS. A two-way ANOVA with the factors treatment (Sham or VNS) and region (PL or IL) found significant effects for both factors (treatment, F_(1,27)_ = 7.670, p = 0.010; region, F_(1,27)_ = 9.98, p = 0.0039), but there was no interaction effect (F_(1,27)_ = 0.5619, p = 0.46; Figure 5F). Post hoc testing with Fisher’s LSD test showed that VNS decreased overall cFos expression in both the PL (p = 0.0492) and in the IL (p = 0.0077). We then examined the activation of PVIs in the PL and IL during cue-induced reinstatement by measuring colocalization of PVI and cFos immunofluorescence. A two-way ANOVA with factors of treatment (Sham or VNS) and region (PL or IL) found no significant effects for both factors, but a significant effect in the interaction of factors (treatment, F_(1,27)_ = 0.5732, p = 0.4556; region, F_(1,27)_ = 0.5152, p = 0.479; interaction, F_(1,27)_ = 17.97, p = 0.0002; Figure 5G). Post hoc testing with Fisher’s LSD test showed that in Sham-treated rats PVI activity was lower in the PL compared to the IL (p = 0.0019), while in VNS-treated rats the PL showed greater cFos activity in PVIs compared to the IL (p = 0.0174). In the PL VNS increased PVI activity (p = 0.0474), but in the IL VNS decreased PVI activity (p =0.0022).

**Figure 5:**
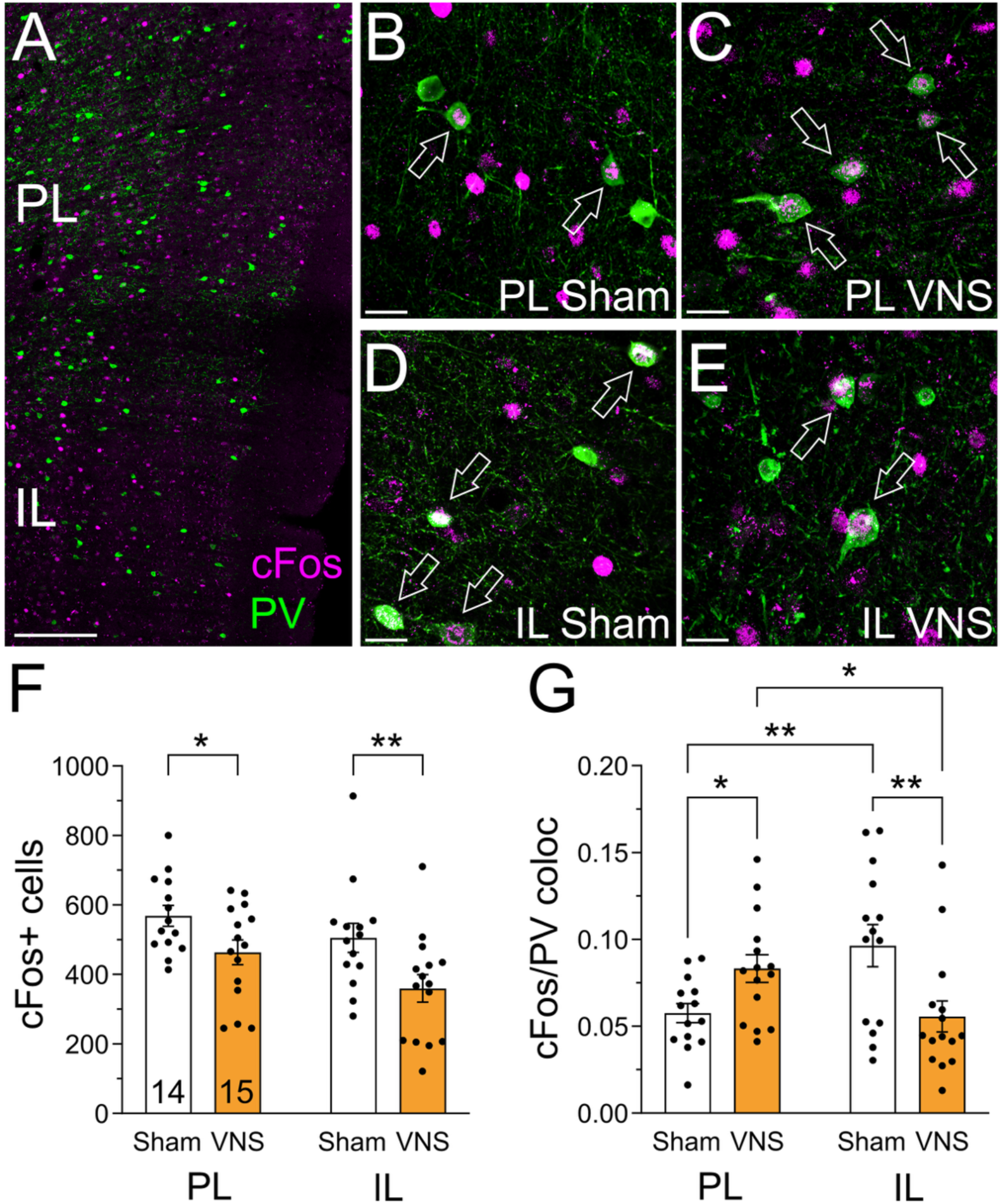
VNS affects activity of parvalbumin-positive interneurons (PVI) in the mPFC during drug-seeking. A) cFos-positive cells (pink) and PVI (green) in the PL and IL. B, C) Examples of cFos+ and parvalbumin+ cells in the PL of Sham- (B), and VNS-stimulated rats (C). D, E) cFos+ and parvalbumin+ cells in the IL of Sham- (D), and VNS-stimulated rats (E). Arrows indicate PVI colocalized with cFos (white). F) VNS decreased overall cFos expression in the PL and IL. G) Colocalization of cFos and PV in the PL and IL. VNS increased the activity of PVI neurons in the PL but decreased cFos activity in PVI of the IL. Scale bars represent 100 µm in A and 20 µm in B-E. *P* values are (*) <0.05, and (**) <0.01.

## Discussion

VNS can modulate cortical and subcortical circuits to improve extinction and reduce drug-seeking (Childs, DeLeon et al. 2017, Driskill, Childs et al. 2024). Here, we examined VNS-induced changes in cellular activity during drug-seeking in two brain areas that project to the IL and PL subregions of the mPFC which are important for drug seeking and the expression of extinction memories. We measured cFos expression in the BLA and vHPC, following cue-induced reinstatement, and we used retrograde labeling of projections to the IL and PL, respectively, to measure pathway-specific activation and modulation by VNS. We found that pairing extinction with VNS reduced reinstatement as previously described (Childs, DeLeon et al. 2017, Driskill, Childs et al. 2024). VNS altered activity in projections from the vHPC and BLA to the mPFC in a pathway-specific manner. In addition, VNS differentially altered activity of PVIs in the IL and PL, which are targets of projections from the BLA and vHPC.

### Reduction of drug seeking by VNS

Exposure to drug-associated cues or stress induces craving and relapse in abstinent patients with substance use disorders (SUD). Extinction forms new associations that compete with these triggers to inhibit drug seeking; however, extinction is often not sufficient to reduce cue-reactivity and prevent relapse (Conklin and Tiffany 2002), potentially because the brain areas required for extinction learning themselves become dysregulated by chronic drug use (Fowler, Volkow et al. 2007, Stephens and Duka 2008, Liu, Liang et al. 2009, Nic Dhonnchadha and Kantak 2011). We replicate previous findings showing that pairing extinction training with VNS accelerates extinction learning and reduces cue-induced reinstatement, suggesting enhanced consolidation of extinction memories (Childs, DeLeon et al. 2017, Driskill, Childs et al. 2024). Vagus nerve stimulation provides a tool to enhance activity-dependent plasticity in neuronal circuits because it leads to the release of multiple neuromodulators (Hassert, Miyashita et al. 2004, Dorr and Debonnel 2006, Roosevelt, Smith et al. 2006, Manta, Dong et al. 2009, Furmaga, Shah et al. 2011, Nichols, Nichols et al. 2011, Manta, El Mansari et al. 2013), that enhance cortical and subcortical synaptic plasticity (Zuo, Smith et al. 2007, Engineer, Riley et al. 2011, Porter, Khodaparast et al. 2012, Pena, Childs et al. 2014, Childs, DeLeon et al. 2017, Hulsey, Shedd et al. 2019, Olsen, Moore et al. 2022). Pairing extinction training with VNS appears to facilitate and consolidate the synaptic plasticity that results from extinction training alone (Childs, DeLeon et al. 2017, Driskill, Childs et al. 2024). BOLD fMRI data in rats also suggests that VNS can reorganize functional connectivity within the limbic system, specifically in circuits that mediate memory and learning, including the amygdala and hippocampus (Cao, Lu et al. 2017, Hachem, Wong et al. 2018).

We have previously shown that VNS-induced reductions in reinstatement correlate with reduced phospho-CREB (pCREB) expression in the BLA and IL (but not the PL) compared with sham-stimulated rats. This overall reduction in pCREB activity in BLA and IL was accompanied by a strengthening of the projection from the IL to the BLA (Childs, DeLeon et al. 2017), suggesting a “focusing” of activity on cells that control the suppression of drug-seeking. Within given brain structures only subpopulations of neurons drive drug-seeking behaviors (Bossert, Stern et al. 2011, Cruz, Babin et al. 2014, Moorman and Aston-Jones 2015, Warren, Kane et al. 2019, Forget, Garcia et al. 2022, Thibeault, Leonard et al. 2025), and the impact of these subpopulations on drug-seeking depends on their projection targets (Augur, Wyckoff et al. 2016, McGlinchey, James et al. 2016, Warren, Kane et al. 2019, Zinsmaier, Dong et al. 2022).

### The role of the mPFC in drug-seeking

To further examine VNS-induced changes in networks that are important for the extinction and cue-induced reinstatement of drug-seeking, we focused on inputs to the mPFC.

Within the mPFC, the PL is often associated with the maintenance of drug-seeking behavior under conditions where cues are predictive of reward. The PL drives reinstatement via its projection to the NAc core. In contrast, the IL has been implicated in the suppression of drug-seeking, by aiding the formation and expression of extinction behaviors (Killcross and Coutureau 2003, Di Ciano, Benham-Hermetz et al. 2007, Peters, LaLumiere et al. 2008, Peters, Vallone et al. 2008, LaLumiere, Niehoff et al. 2010, Van den Oever, Rotaru et al. 2013, Augur, Wyckoff et al. 2016, Gutman, Nett et al. 2017, Muller Ewald, De Corte et al. 2019, Nett, Zimbelman et al. 2023). However, because frontal cortical areas are strongly interconnected, information about actions, emotions, and stimuli is available to all prefrontal regions (Euston, Gruber et al. 2012, Anastasiades and Carter 2021) and a number of studies have indicated overlapping roles for the PL and IL in reward seeking (Moorman and Aston-Jones 2015, Gentry and Roesch 2018, Madangopal, Ramsey et al. 2021), as reviewed in (Howland, Ito et al. 2022).

Here, we focused on inputs from the BLA and vHPC that are important in appetitive associative learning and contextual processing. The goal was to determine whether neurons from these areas that send direct projections to the PL or IL, respectively, are selectively activated during reinstatement and whether pairing extinction training with VNS alters these activation patterns.

### VNS modulation of BLA projections to the mPFC

The BLA is essential for cue-induced cocaine-seeking and relapse, serving as a critical hub that processes drug-associated memories and drives cocaine-seeking behavior, particularly in response to drug-associated cues. Furthermore, the BLA plays a complex dual role in extinction learning, being necessary for both the formation of cocaine-cue associations and their subsequent extinction (Di Ciano and Everitt 2004, Fuchs, Feltenstein et al. 2006, Stefanik and Kalivas 2013, Pelloux, Minier-Toribio et al. 2018). We have previously shown that VNS-induced suppression of cue-induced reinstatement following extinction correlates with a reduced expression of phospho-CREB in the BLA (Childs, DeLeon et al. 2017). Here we observed a similar global reduction in cFos signal in the BLA of VNS-treated rats, suggesting that VNS overall reduces activity in BLA networks. This reduction in activity may reflect weakened cue-reward associations (Fuchs, Feltenstein et al. 2006, Fuchs, Bell et al. 2009) or reduced motivational drive when exposed to triggering stimuli (Servonnet, Hernandez et al. 2020).

The BLA-mPFC pathway is integral in assigning emotional salience to drug-associated cues, facilitating the formation of strong drug-context associations that sustain addictive behaviors (Kantak, Black et al. 2002, Kalivas and McFarland 2003, McLaughlin and See 2003, Di Ciano and Everitt 2004, Peters, Vallone et al. 2008, Stefanik and Kalivas 2013). Importantly, previous studies have shown that PL-innervating and IL-innervating neurons in BLA have different molecular markers and functions, with PL-innervating neurons encoding predominantly negative valence, while IL-projecting neurons mediate positive behaviors (Kim, Pignatelli et al. 2016). Importantly, intermingled subpopulations of BLA projection neurons that target the PL and the IL, respectively, are differentially recruited during either the expression of fear (PL) or during fear extinction (IL) (Senn, Wolff et al. 2014). We hypothesized that the BLA regulates mPFC activity in a similar manner during the extinction of drug-seeking and that VNS would enhance activity in the IL-projecting BLA neurons to facilitate or consolidate extinction to reduce responding during cue-induced reinstatement. Alternatively, VNS might weaken inputs from the BLA to the PL which drive drug-seeking (Stefanik and Kalivas 2013). Surprisingly, we observed the opposite activation pattern in VNS-treated animals: PL-projecting neurons showed more cFos activity after cue-induced reinstatement, while IL-projecting neurons in the BLA showed less cFos activity. However, the overall influence of glutamatergic inputs from the BLA to the mPFC is predominantly inhibitory: BLA inputs make their strongest connections onto parvalbumin- and somatostatin-expressing interneurons, which provide robust feedforward inhibition onto IL projection neurons, including cortico-amygdala neurons that project back to the BLA (Gabbott, Warner et al. 2006, Dilgen, Tejeda et al. 2013, McGarry and Carter 2016). Thus, the VNS-induced reduction of activity in BLA◊IL neurons is likely to disinhibit the IL, aiding expression of extinction memories. Similarly, enhanced feedforward inhibition in the BLA◊PL projection following VNS during extinction training might reduce PL activity and drug-seeking. Taken together, the opposing VNS modulation of BLA pathways may facilitate IL-dependent extinction mechanisms while suppressing PL-dependent drug-seeking processes.

### VNS modulation of vHPC projections to the mPFC

The hippocampus is involved in the integration of contextual information, emotional states, and memory processes, which in turn can trigger craving and relapse when reactivated (Neisewander, Baker et al. 2000, Ferbinteanu and McDonald 2001, Kalivas and McFarland 2003, Sun and Rebec 2003, Rogers and See 2007, Atkins, Mashhoon et al. 2008, Lasseter, Xie et al. 2010, Goode and Maren 2019). Direct inputs from the hippocampus to the mPFC originate mainly from the vHPC (Jay and Witter 1991, Cenquizca and Swanson 2007, Aleman-Andrade, Witter et al. 2025). Electrical stimulation of the ventral subiculum, which serves as output region of the vHPC, elicits reinstatement of extinguished drug-seeking behaviors (Vorel, Liu et al. 2001, Taepavarapruk and Phillips 2003). In contrast, inactivation of either the vHPC or the ventral subiculum impairs both cue-induced and cocaine-primed cocaine-seeking behaviors (Sun and Rebec 2003, Rogers and See 2007), demonstrating their critical role in both the formation and retrieval of drug-associated memories.

The vHPC-mPFC pathway is integral to the regulation of behaviors that require the integration of contextual and emotional information with higher-order cognitive processes (Vertes 2006) and as such the projections from the vHPC to the mPFC are important for contextual processing and memory retrieval in drug-seeking and relapse. Specifically, renewal of heroin-seeking has been shown to engage the vHPC◊IL circuitry (Bossert, Adhikary et al. 2016, Wang, Ge et al. 2018), and inactivation of vHPC◊IL neurons prevents context-induced reinstatement of heroin seeking (Wang, Ge et al. 2018). In contrast, the vHPC◊PL pathway does not appear necessary for the reinstatement of drug-seeking (Bossert, Adhikary et al. 2016, Wang, Ge et al. 2018).

In addition, vHPC inputs to the mPFC also aid the consolidation and retrieval of extinction memories, thus contributing to the attenuation of conditioned responses to drug-related stimuli (Castilla-Ortega, Serrano et al. 2016, Kutlu and Gould 2016). Importantly, the vHPC-mPFC circuitry is bidirectional, which enables the dynamic regulation of behavior in response to changing environmental contexts, as is the case during extinction learning. These functions of the vHPC, and the vHPC-mPFC projection specifically, may render it a prominent target for VNS modulation of synaptic plasticity during extinction learning. In our data we found no differences between VNS- and Sham-stimulated rats in the total number of cFos+ cells in the vHPC, indicating that pairing extinction with VNS did not affect overall activity of the vHPC during reinstatement. However, we observed pathway-specific alterations in the number of cFos+ vHPC cells, such that in VNS-treated rats IL-projecting cells (but not cells projecting to the PL) showed a selective reduction of cFos expression. Previous work has shown that activation of the vHPC◊IL pathway engages strong inhibition of IL outputs, preventing extinction mechanisms of both drug-seeking (Bossert, Adhikary et al. 2016, Wang, Ge et al. 2018) and fear renewal (Marek, Jin et al. 2018). Thus, the VNS-induced reduction of activity in this pathway may result in disinhibition and may facilitate the expression of extinction memories from the IL. In contrast, the lack of VNS modulation of the vHPC◊PL projection could either further support a functional dissociation between vHPC projections to the IL and PL as previously suggested (Bossert, Adhikary et al. 2016, Wang, Ge et al. 2018), or it may reflect the relatively weaker innervation from the vHPC to the PL (Wang, Jin et al. 2016, Goode and Maren 2019).

### VNS modulation of PVI in the mPFC

Afferents from both the vHPC and the BLA directly contact pyramidal cells and interneurons in the mPFC. As outlined above projections onto PVI and somatostatin-expressing interneurons have been shown to exert strong inhibitory effects on the functions of the PL and IL (Tierney, Degenetais et al. 2004, McGarry and Carter 2016, Abbas, Sundiang et al. 2018, Liu and Carter 2018, Marek, Jin et al. 2018, Sun, Li et al. 2019, Yang, Mack et al. 2021, Sanchez-Bellot, AlSubaie et al. 2022, Aleman-Andrade, Witter et al. 2025). Importantly, PVI in the PL and IL receive inputs from largely different neurons within the same brain areas (Sun, Li et al. 2019). Activation of PL PVI supports the transition from a cue-driven to a flexible behavioral state, thus facilitating extinction of reward-seeking behavior (Sparta, Hovelso et al. 2014). We examined VNS-induced changes in cFos expression in the mPFC, as well as cFos colocalization with PVI in the PL and IL to determine if VNS may alter PVI activity to facilitate extinction and reduce reinstatement. Consistent with our previous findings using the activitiy marker pCREB (Childs, DeLeon et al. 2017), we found an overall reduction in cFos activity in both PL and IL. Thus while cFos expression can serve as a general marker for VNS-induced changes in activity in the mPFC, it is by itself not sufficient to distinguish potentiall different roles of multiple overlapping networks for drug-seeking or extinction in the IL and PL. However, when we examined the colocalization of cFos with PVI we found that VNS caused opposing, area-specific changes: PVI in the PL showed increased cFos activity, which may reflect increased inhibition in the PL and may cause reduced drug-seeking. This increase in cFos expression is also consistent with an enhanced drive from upstream BLA in VNS animals (Figure 3J). In contrast, PVI in the IL showed less cFos activity, indicating VNS-induced disinhibition of IL networks. This is also consistent with our finding that VNS reduced activity in the projections from the vHPC and BLA, and previous findings outlined above which show that inactivation of IL-projecting neurons prevents reinstatement of fear and drug-seeking (Marek, Jin et al. 2018, Wang, Ge et al. 2018, Binette, Liu et al. 2023). Taken together, VNS may reduce drug-seeking by inhibiting activity in the PL and also causing disinhibition of the IL by reducing activity in IL PVI, enabling expression of extinction memories. These effects may be driven by VNS-induced changes in projections from the vHPC and the BLA. Our findings have important limitations: Our results are correlational and do not establish causal relationships between VNS-induced changes in PVI activity, alterations in upstream projections to the mPFC, and behavioral outcomes. Additionally, projections from the vHPC and BLA innervate other interneuron populations beyond PVIs, including somatostatin and vasoactive intestinal peptide interneurons (McGarry and Carter 2016, Sun, Li et al. 2019). These interneuron subtypes powerfully regulate mPFC function and can themselves inhibit PVIs (Xu, Liu et al. 2019, Yang, Mack et al. 2021). Therefore, future studies must establish direct causal links between VNS-induced changes in mPFC projections and the activity of PVIs and other interneuron populations in regulating drug-seeking behavior. Furthermore, our study only used male rats, thus sex differences in the circuits through which VNS modulates extinction learning cannot be excluded. However, we have recently shown that VNS reduces cue-induced reinstatement and modulates plasticity in the BLA to IL circuit in both male and female rats in the same way (Arezoomandan, Vu et al. 2025).

## Summary

Our work demonstrates that vagus nerve stimulation (VNS) paired with extinction training reduces drug-seeking behavior by selectively modulating prefrontal cortex circuits. We found that VNS differentially affects parvalbumin interneurons: increasing inhibitory activity in the prelimbic cortex (which drives drug-seeking) while decreasing it in the infralimbic cortex (which supports extinction). VNS also altered pathway-specific projections from the ventral hippocampus and basolateral amygdala, creating enhanced extinction circuits while suppressing relapse-driving networks. Our findings reveal how neuromodulation can precisely target competing neural circuits in addiction, suggesting VNS could serve as an adjunctive therapy for substance use disorders by strengthening the brain’s natural recovery mechanisms while weakening cue-driven relapse pathways.

## Acknowledgements

Cocaine HCl was provided by the NIDA Drug Supply Program. The content is solely the responsibility of the authors and does not necessarily represent the official views of the National Institute on Drug Abuse or the National Institutes of Health.

## Author Contributions

CD, LV Conceptualization, data acquisition and analysis, statistical analysis, writing, and editing the manuscript. **SJ, FS, LW, NS, ST, AK, ZH, RN, ZK, NM**: Data acquisition and analysis. **SK**: Funding acquisition, conceptualization, writing, and editing the manuscript. All authors reviewed the manuscript and approved the final version for publication.

## Data availability statement

The data that support the findings of this study are available from the corresponding author upon reasonable request.

## Ethical approval

This study has been conducted based on ethical approval from the University of Texas at Dallas.

## References

1. Abbas, A. I., M. J. M. Sundiang, B. Henoch, M. P. Morton, S. S. Bolkan, A. J. Park, A. Z. Harris, C. Kellendonk and J. A. Gordon (2018). “Somatostatin Interneurons Facilitate Hippocampal-Prefrontal Synchrony and Prefrontal Spatial Encoding.” Neuron 100(4): 926–939 e923.

2. Aleman-Andrade, P., M. P. Witter, K. I. Tsutsui and S. Ohara (2025). “Dorsal-caudal and ventral hippocampus target different cell populations in the medial frontal cortex in rodents.” J Neurosci.

3. Anastasiades, P. G. and A. G. Carter (2021). “Circuit organization of the rodent medial prefrontal cortex.” Trends Neurosci 44(7): 550–563.

4. Arezoomandan, R., L. Vu, C. Driskill and S. Kroener (2025). “Vagus nerve stimulation modulates synaptic plasticity induced by cocaine- seeking in reward-related circuitry.” Res Sq.

5. Atkins, A. L., Y. Mashhoon and K. M. Kantak (2008). “Hippocampal regulation of contextual cue-induced reinstatement of cocaine-seeking behavior.” Pharmacol Biochem Behav 90(3): 481–491.

6. Augur, I. F., A. R. Wyckoff, G. Aston-Jones, P. W. Kalivas and J. Peters (2016). “Chemogenetic Activation of an Extinction Neural Circuit Reduces Cue-Induced Reinstatement of Cocaine Seeking.” J Neurosci 36(39): 10174–10180.

7. Binette, A. N., J. Liu, H. Bayer, K. L. Crayton, L. Melissari, S. O. Sweck and S. Maren (2023). “Parvalbumin-Positive Interneurons in the Medial Prefrontal Cortex Regulate Stress-Induced Fear Extinction Impairments in Male and Female Rats.” J Neurosci 43(22): 4162–4173.

8. Bossert, J. M., S. Adhikary, R. St Laurent, N. J. Marchant, H. L. Wang, M. Morales and Y. Shaham (2016). “Role of projections from ventral subiculum to nucleus accumbens shell in context-induced reinstatement of heroin seeking in rats.” Psychopharmacology (Berl) 233(10): 1991–2004.

9. Bossert, J. M., A. L. Stern, F. R. Theberge, C. Cifani, E. Koya, B. T. Hope and Y. Shaham (2011). “Ventral medial prefrontal cortex neuronal ensembles mediate context-induced relapse to heroin.” Nat Neurosci 14(4): 420–422.

10. Cao, B., J. Wang, M. Shahed, B. Jelfs, R. H. Chan and Y. Li (2016). “Vagus Nerve Stimulation Alters Phase Synchrony of the Anterior Cingulate Cortex and Facilitates Decision Making in Rats.” Sci Rep 6: 35135.

11. Cao, J., K. H. Lu, T. L. Powley and Z. Liu (2017). “Vagal nerve stimulation triggers widespread responses and alters large-scale functional connectivity in the rat brain.” PLoS One 12(12): e0189518.

12. Castilla-Ortega, E., A. Serrano, E. Blanco, P. Araos, J. Suarez, F. J. Pavon, F. Rodriguez de Fonseca and L. J. Santin (2016). “A place for the hippocampus in the cocaine addiction circuit: Potential roles for adult hippocampal neurogenesis.” Neurosci Biobehav Rev 66: 15–32.

13. Cenquizca, L. A. and L. W. Swanson (2007). “Spatial organization of direct hippocampal field CA1 axonal projections to the rest of the cerebral cortex.” Brain Res Rev 56(1): 1–26.

14. Childs, J. E., A. C. Alvarez-Dieppa, C. K. McIntyre and S. Kroener (2015). “Vagus Nerve Stimulation as a Tool to Induce Plasticity in Pathways Relevant for Extinction Learning.” J Vis Exp(102): e53032.

15. Childs, J. E., J. DeLeon, E. Nickel and S. Kroener (2017). “Vagus nerve stimulation reduces cocaine seeking and alters plasticity in the extinction network.” Learn Mem 24(1): 35–42.

16. Childs, J. E., J. DeLeon, E. Nickel and S. Kroener (2017). “Vagus nerve stimulation reduces cocaine seeking and alters plasticity in the extinction network.” Learning & Memory 24(1): 3542.

17. Childs, J. E., S. Kim, C. M. Driskill, E. Hsiu and S. Kroener (2019). “Vagus nerve stimulation during extinction learning reduces conditioned place preference and context-induced reinstatement of cocaine seeking.” Brain Stimul 12(6): 1448–1455.

18. Clark, K. B., D. K. Naritoku, D. C. Smith, R. A. Browning and R. A. Jensen (1999). “Enhanced recognition memory following vagus nerve stimulation in human subjects.” Nat Neurosci 2(1): 9498.

19. Clark, K. B., D. C. Smith, D. L. Hassert, R. A. Browning, D. K. Naritoku and R. A. Jensen (1998). “Posttraining electrical stimulation of vagal afferents with concomitant vagal efferent inactivation enhances memory storage processes in the rat.” Neurobiol Learn Mem 70(3): 364–373.

20. Conklin, C. A. and S. T. Tiffany (2002). “Applying extinction research and theory to cue-exposure addiction treatments.” Addiction 97(2): 155–167.

21. Cruz, F. C., K. R. Babin, R. M. Leao, E. M. Goldart, J. M. Bossert, Y. Shaham and B. T. Hope (2014). “Role of nucleus accumbens shell neuronal ensembles in context-induced reinstatement of cocaine-seeking.” J Neurosci 34(22): 7437–7446.

22. Cruz, F. C., F. Javier Rubio and B. T. Hope (2015). “Using c-fos to study neuronal ensembles in corticostriatal circuitry of addiction.” Brain Res 1628(Pt A): 157–173.

23. Di Ciano, P., J. Benham-Hermetz, A. P. Fogg and G. E. Osborne (2007). “Role of the prelimbic cortex in the acquisition, re-acquisition or persistence of responding for a drug-paired conditioned reinforcer.” Neuroscience 150(2): 291–298.

24. Di Ciano, P. and B. J. Everitt (2004). “Direct interactions between the basolateral amygdala and nucleus accumbens core underlie cocaine-seeking behavior by rats.” J Neurosci 24(32): 7167-7173.

25. Dilgen, J., H. A. Tejeda and P. O’Donnell (2013). “Amygdala inputs drive feedforward inhibition in the medial prefrontal cortex.” J Neurophysiol 110(1): 221–229.

26. Dorr, A. E. and G. Debonnel (2006). “Effect of vagus nerve stimulation on serotonergic and noradrenergic transmission.” J Pharmacol Exp Ther 318(2): 890–898.

27. Driskill, C. M., J. E. Childs, B. Itmer, J. S. Rajput and S. Kroener (2022). “Acute Vagus Nerve Stimulation Facilitates Short Term Memory and Cognitive Flexibility in Rats.” Brain Sci 12(9).

28. Driskill, C. M., J. E. Childs, A. J. Phensy, S. R. Rodriguez, J. T. O’Brien, K. L. Lindquist, A. Naderi, B. Bordieanu, J. F. McGinty and S. Kroener (2024). “Vagus Nerve Stimulation (VNS) Modulates Synaptic Plasticity in the Infralimbic Cortex via Trk-B Receptor Activation to Reduce Drug-Seeking in Male Rats.” J Neurosci 44(23).

29. Engineer, N. D., J. R. Riley, J. D. Seale, W. A. Vrana, J. A. Shetake, S. P. Sudanagunta, M. S. Borland and M. P. Kilgard (2011). “Reversing pathological neural activity using targeted plasticity.” Nature 470(7332): 101–104.

30. Euston, D. R., A. J. Gruber and B. L. McNaughton (2012). “The role of medial prefrontal cortex in memory and decision making.” Neuron 76(6): 1057–1070.

31. Ferbinteanu, J. and R. J. McDonald (2001). “Dorsal/ventral hippocampus, fornix, and conditioned place preference.” Hippocampus 11(2): 187–200.

32. Forget, B., E. M. Garcia, A. Godino, L. D. Rodriguez, V. Kappes, P. Poirier, A. Andrianarivelo, E. S. Marchan, M. C. Allichon, M. Marias, P. Vanhoutte, J. A. Girault, R. Maldonado and J. Caboche (2022). “Cell-type- and region-specific modulation of cocaine seeking by micro-RNA-1 in striatal projection neurons.” Mol Psychiatry 27(2): 918–928.

33. Fowler, J. S., N. D. Volkow, C. A. Kassed and L. Chang (2007). “Imaging the addicted human brain.” Sci Pract Perspect 3(2): 4–16.

34. Fuchs, R. A., G. H. Bell, D. R. Ramirez, J. L. Eaddy and Z. I. Su (2009). “Basolateral amygdala involvement in memory reconsolidation processes that facilitate drug context-induced cocaine seeking.” Eur J Neurosci 30(5): 889–900.

35. Fuchs, R. A., M. W. Feltenstein and R. E. See (2006). “The role of the basolateral amygdala in stimulus-reward memory and extinction memory consolidation and in subsequent conditioned cued reinstatement of cocaine seeking.” Eur J Neurosci 23(10): 2809–2813.

36. Furmaga, H., A. Shah and A. Frazer (2011). “Serotonergic and noradrenergic pathways are required for the anxiolytic-like and antidepressant-like behavioral effects of repeated vagal nerve stimulation in rats.” Biol Psychiatry 70(10): 937–945.

37. Gabbott, P. L., T. A. Warner and S. J. Busby (2006). “Amygdala input monosynaptically innervates parvalbumin immunoreactive local circuit neurons in rat medial prefrontal cortex.” Neuroscience 139(3): 1039–1048.

38. Gentry, R. N. and M. R. Roesch (2018). “Neural Activity in Ventral Medial Prefrontal Cortex Is Modulated More Before Approach Than Avoidance During Reinforced and Extinction Trial Blocks.” J Neurosci 38(19): 4584–4597.

39. Goode, T. D. and S. Maren (2019). “Common neurocircuitry mediating drug and fear relapse in preclinical models.” Psychopharmacology (Berl) 236(1): 415–437.

40. Gourley, S. L. and J. R. Taylor (2016). “Going and stopping: Dichotomies in behavioral control by the prefrontal cortex.” Nat Neurosci 19(6): 656–664.

41. Gutman, A. L., K. E. Nett, C. V. Cosme, W. R. Worth, S. C. Gupta, J. A. Wemmie and R. T. LaLumiere (2017). “Extinction of Cocaine Seeking Requires a Window of Infralimbic Pyramidal Neuron Activity after Unreinforced Lever Presses.” J Neurosci 37(25): 6075–6086.

42. Hachem, L. D., S. M. Wong and G. M. Ibrahim (2018). “The vagus afferent network: emerging role in translational connectomics.” Neurosurg Focus 45(3): E2.

43. Hassert, D. L., T. Miyashita and C. L. Williams (2004). “The effects of peripheral vagal nerve stimulation at a memory-modulating intensity on norepinephrine output in the basolateral amygdala.” Behav Neurosci 118(1): 79–88.

44. Havermans, R. C. and A. T. Jansen (2003). “Increasing the efficacy of cue exposure treatment in preventing relapse of addictive behavior.” Addict Behav 28(5): 989–994.

45. Howland, J. G., R. Ito, C. C. Lapish and F. R. Villaruel (2022). “The rodent medial prefrontal cortex and associated circuits in orchestrating adaptive behavior under variable demands.” Neurosci Biobehav Rev 135: 104569.

46. Hulsey, D. R., C. M. Shedd, S. F. Sarker, M. P. Kilgard and S. A. Hays (2019). “Norepinephrine and serotonin are required for vagus nerve stimulation directed cortical plasticity.” Exp Neurol 320: 112975.

47. Jay, T. M. and M. P. Witter (1991). “Distribution of hippocampal CA1 and subicular efferents in the prefrontal cortex of the rat studied by means of anterograde transport of Phaseolus vulgaris-leucoagglutinin.” J Comp Neurol 313(4): 574–586.

48. Kalivas, P. W. and K. McFarland (2003). “Brain circuitry and the reinstatement of cocaine-seeking behavior.” Psychopharmacology (Berl) 168(1-2): 44–56.

49. Kantak, K. M., Y. Black, E. Valencia, K. Green-Jordan and H. B. Eichenbaum (2002). “Dissociable effects of lidocaine inactivation of the rostral and caudal basolateral amygdala on the maintenance and reinstatement of cocaine-seeking behavior in rats.” J Neurosci 22(3): 1126–1136.

50. Killcross, S. and E. Coutureau (2003). “Coordination of actions and habits in the medial prefrontal cortex of rats.” Cereb Cortex 13(4): 400–408.

51. Kim, J., M. Pignatelli, S. Xu, S. Itohara and S. Tonegawa (2016). “Antagonistic negative and positive neurons of the basolateral amygdala.” Nat Neurosci 19(12): 1636–1646.

52. Krahl, S. E. and K. B. Clark (2012). “Vagus nerve stimulation for epilepsy: A review of central mechanisms.” Surg Neurol Int 3(Suppl 4): S255–259.

53. Kutlu, M. G. and T. J. Gould (2016). “Effects of drugs of abuse on hippocampal plasticity and hippocampus-dependent learning and memory: contributions to development and maintenance of addiction.” Learn Mem 23(10): 515–533.

54. LaLumiere, R. T., K. E. Niehoff and P. W. Kalivas (2010). “The infralimbic cortex regulates the consolidation of extinction after cocaine self-administration.” Learn Mem 17(4): 168–175.

55. Lasseter, H. C., X. Xie, D. R. Ramirez and R. A. Fuchs (2010). “Sub-region specific contribution of the ventral hippocampus to drug context-induced reinstatement of cocaine-seeking behavior in rats.” Neuroscience 171(3): 830–839.

56. Liu, J., J. Liang, W. Qin, J. Tian, K. Yuan, L. Bai, Y. Zhang, W. Wang, Y. Wang, Q. Li, L. Zhao, L. Lu, K. M. von Deneen, Y. Liu and M. S. Gold (2009). “Dysfunctional connectivity patterns in chronic heroin users: an fMRI study.” Neurosci Lett 460(1): 72–77.

57. Liu, X. and A. G. Carter (2018). “Ventral Hippocampal Inputs Preferentially Drive Corticocortical Neurons in the Infralimbic Prefrontal Cortex.” J Neurosci 38(33): 7351–7363.

58. Madangopal, R., L. A. Ramsey, S. J. Weber, M. B. Brenner, V. A. Lennon, O. R. Drake, L. E. Komer, B. J. Tunstall, J. M. Bossert, Y. Shaham and B. T. Hope (2021). “Inactivation of the infralimbic cortex decreases discriminative stimulus-controlled relapse to cocaine seeking in rats.” Neuropsychopharmacology 46(11): 1969–1980.

59. Manta, S., J. Dong, G. Debonnel and P. Blier (2009). “Enhancement of the function of rat serotonin and norepinephrine neurons by sustained vagus nerve stimulation.” J Psychiatry Neurosci 34(4): 272–280.

60. Manta, S., M. El Mansari, G. Debonnel and P. Blier (2013). “Electrophysiological and neurochemical effects of long-term vagus nerve stimulation on the rat monoaminergic systems.” Int J Neuropsychopharmacol 16(2): 459–470.

61. Marek, R., J. Jin, T. D. Goode, T. F. Giustino, Q. Wang, G. M. Acca, R. Holehonnur, J. E. Ploski, P. J. Fitzgerald, T. Lynagh, J. W. Lynch, S. Maren and P. Sah (2018). “Hippocampus-driven feed-forward inhibition of the prefrontal cortex mediates relapse of extinguished fear.” Nat Neurosci 21(3): 384–392.

62. McGarry, L. M. and A. G. Carter (2016). “Inhibitory Gating of Basolateral Amygdala Inputs to the Prefrontal Cortex.” J Neurosci 36(36): 9391–9406.

63. McGlinchey, E. M., M. H. James, S. V. Mahler, C. Pantazis and G. Aston-Jones (2016). “Prelimbic to Accumbens Core Pathway Is Recruited in a Dopamine-Dependent Manner to Drive Cued Reinstatement of Cocaine Seeking.” J Neurosci 36(33): 8700–8711.

64. McLaughlin, J. and R. E. See (2003). “Selective inactivation of the dorsomedial prefrontal cortex and the basolateral amygdala attenuates conditioned-cued reinstatement of extinguished cocaine-seeking behavior in rats.” Psychopharmacology (Berl) 168(1-2): 57–65.

65. Millan, E. Z., N. J. Marchant and G. P. McNally (2011). “Extinction of drug seeking.” Behav Brain Res 217(2): 454–462.

66. Moorman, D. E. and G. Aston-Jones (2015). “Prefrontal neurons encode context-based response execution and inhibition in reward seeking and extinction.” Proc Natl Acad Sci U S A 112(30): 9472–9477.

67. Muller Ewald, V. A., B. J. De Corte, S. C. Gupta, K. V. Lillis, N. S. Narayanan, J. A. Wemmie and R. T. LaLumiere (2019). “Attenuation of cocaine seeking in rats via enhancement of infralimbic cortical activity using stable step-function opsins.” Psychopharmacology (Berl) 236(1): 479–490.

68. Neisewander, J. L., D. A. Baker, R. A. Fuchs, L. T. Tran-Nguyen, A. Palmer and J. F. Marshall (2000). “Fos protein expression and cocaine-seeking behavior in rats after exposure to a cocaine self-administration environment.” J Neurosci 20(2): 798–805.

69. Nemeroff, C. B., H. S. Mayberg, S. E. Krahl, J. McNamara, A. Frazer, T. R. Henry, M. S. George, D. S. Charney and S. K. Brannan (2006). “VNS therapy in treatment-resistant depression: clinical evidence and putative neurobiological mechanisms.” Neuropsychopharmacology 31(7): 1345-1355.

70. Nett, K. E., A. R. Zimbelman, M. S. McGregor, V. Alizo Vera, M. R. Harris and R. T. LaLumiere (2023). “Infralimbic Projections to the Nucleus Accumbens Shell and Amygdala Regulate the Encoding of Cocaine Extinction Learning.” J Neurosci 43(8): 1348–1359.

71. Nic Dhonnchadha, B. A. and K. M. Kantak (2011). “Cognitive enhancers for facilitating drug cue extinction: insights from animal models.” Pharmacol Biochem Behav 99(2): 229–244.

72. Nichols, J. A., A. R. Nichols, S. M. Smirnakis, N. D. Engineer, M. P. Kilgard and M. Atzori (2011). “Vagus nerve stimulation modulates cortical synchrony and excitability through the activation of muscarinic receptors.” Neuroscience 189: 207–214.

73. O’Brien, C. P., A. R. Childress, R. Ehrman and S. J. Robbins (1998). “Conditioning factors in drug abuse: can they explain compulsion?” J Psychopharmacol 12(1): 15–22.

74. Olsen, L. K., R. J. Moore, N. A. Bechmann, V. T. Ethridge, N. M. Gargas, S. D. Cunningham, Z. Kuang, J. K. Whicker, J. G. Rohan and C. N. Hatcher-Solis (2022). “Vagus nerve stimulation-induced cognitive enhancement: Hippocampal neuroplasticity in healthy male rats.” Brain Stimul 15(5): 1101–1110.

75. Paxinos, G., Watson, C. (2007). The rat brain in stereotaxic coordinates. Amsterdam, Academic Press.

76. Pelloux, Y., A. Minier-Toribio, J. K. Hoots, J. M. Bossert and Y. Shaham (2018). “Opposite Effects of Basolateral Amygdala Inactivation on Context-Induced Relapse to Cocaine Seeking after Extinction versus Punishment.” J Neurosci 38(1): 51–59.

77. Pena, D. F., J. E. Childs, S. Willett, A. Vital, C. K. McIntyre and S. Kroener (2014). “Vagus nerve stimulation enhances extinction of conditioned fear and modulates plasticity in the pathway from the ventromedial prefrontal cortex to the amygdala.” Front Behav Neurosci 8: 327.

78. Perry, C. J. and A. J. Lawrence (2017). “Addiction, cognitive decline and therapy: seeking ways to escape a vicious cycle.” Genes Brain Behav 16(1): 205–218.

79. Peters, J., P. W. Kalivas and G. J. Quirk (2009). “Extinction circuits for fear and addiction overlap in prefrontal cortex.” Learn Mem 16(5): 279–288.

80. Peters, J., R. T. LaLumiere and P. W. Kalivas (2008). “Infralimbic prefrontal cortex is responsible for inhibiting cocaine seeking in extinguished rats.” J Neurosci 28(23): 6046–6053.

81. Peters, J., T. Pattij and T. J. De Vries (2013). “Targeting cocaine versus heroin memories: divergent roles within ventromedial prefrontal cortex.” Trends Pharmacol Sci 34(12): 689–695.

82. Peters, J., J. Vallone, K. Laurendi and P. W. Kalivas (2008). “Opposing roles for the ventral prefrontal cortex and the basolateral amygdala on the spontaneous recovery of cocaine-seeking in rats.” Psychopharmacology (Berl) 197(2): 319–326.

83. Phillips, K. A., D. H. Epstein and K. L. Preston (2014). “Psychostimulant addiction treatment.” Neuropharmacology 87: 150–160.

84. Porter, B. A., N. Khodaparast, T. Fayyaz, R. J. Cheung, S. S. Ahmed, W. A. Vrana, R. L. Rennaker, 2nd and M. P. Kilgard (2012). “Repeatedly pairing vagus nerve stimulation with a movement reorganizes primary motor cortex.” Cereb Cortex 22(10): 2365–2374.

85. Rogers, J. L. and R. E. See (2007). “Selective inactivation of the ventral hippocampus attenuates cue-induced and cocaine-primed reinstatement of drug-seeking in rats.” Neurobiol Learn Mem 87(4): 688–692.

86. Roosevelt, R. W., D. C. Smith, R. W. Clough, R. A. Jensen and R. A. Browning (2006). “Increased extracellular concentrations of norepinephrine in cortex and hippocampus following vagus nerve stimulation in the rat.” Brain Res 1119(1): 124–132.

87. Rush, A. J., L. B. Marangell, H. A. Sackeim, M. S. George, S. K. Brannan, S. M. Davis, R. Howland, M. A. Kling, B. R. Rittberg, W. J. Burke, M. H. Rapaport, J. Zajecka, A. A. Nierenberg, M. M. Husain, D. Ginsberg and R. G. Cooke (2005). “Vagus nerve stimulation for treatment-resistant depression: a randomized, controlled acute phase trial.” Biol Psychiatry 58(5): 347–354.

88. Sanchez-Bellot, C., R. AlSubaie, K. Mishchanchuk, R. W. S. Wee and A. F. MacAskill (2022). “Two opposing hippocampus to prefrontal cortex pathways for the control of approach and avoidance behaviour.” Nat Commun 13(1): 339.

89. Senn, V., S. B. Wolff, C. Herry, F. Grenier, I. Ehrlich, J. Grundemann, J. P. Fadok, C. Muller, J. J. Letzkus and A. Luthi (2014). “Long-range connectivity defines behavioral specificity of amygdala neurons.” Neuron 81(2): 428–437.

90. Servonnet, A., G. Hernandez, C. El Hage, P. P. Rompre and A. N. Samaha (2020). “Optogenetic Activation of the Basolateral Amygdala Promotes Both Appetitive Conditioning and the Instrumental Pursuit of Reward Cues.” J Neurosci 40(8): 1732–1743.

91. Sinha, R., T. Fuse, L. R. Aubin and S. S. O’Malley (2000). “Psychological stress, drug-related cues and cocaine craving.” Psychopharmacology (Berl) 152(2): 140–148.

92. Sparta, D. R., N. Hovelso, A. O. Mason, P. A. Kantak, R. L. Ung, H. K. Decot and G. D. Stuber (2014). “Activation of prefrontal cortical parvalbumin interneurons facilitates extinction of reward-seeking behavior.” J Neurosci 34(10): 3699–3705.

93. Stefanik, M. T. and P. W. Kalivas (2013). “Optogenetic dissection of basolateral amygdala projections during cue-induced reinstatement of cocaine seeking.” Front Behav Neurosci 7: 213.

94. Stephens, D. N. and T. Duka (2008). “Review. Cognitive and emotional consequences of binge drinking: role of amygdala and prefrontal cortex.” Philos Trans R Soc Lond B Biol Sci 363(1507): 3169–3179.

95. Sun, L., J. Perakyla, K. Holm, J. Haapasalo, K. Lehtimaki, K. H. Ogawa, J. Peltola and K. M. Hartikainen (2017). “Vagus nerve stimulation improves working memory performance.” J Clin Exp Neuropsychol 39(10): 954–964.

96. Sun, Q., X. Li, M. Ren, M. Zhao, Q. Zhong, Y. Ren, P. Luo, H. Ni, X. Zhang, C. Zhang, J. Yuan, A. Li, M. Luo, H. Gong and Q. Luo (2019). “A whole-brain map of long-range inputs to GABAergic interneurons in the mouse medial prefrontal cortex.” Nat Neurosci 22(8): 1357–1370.

97. Sun, W. and G. V. Rebec (2003). “Lidocaine inactivation of ventral subiculum attenuates cocaine-seeking behavior in rats.” J Neurosci 23(32): 10258–10264.

98. Taepavarapruk, P. and A. G. Phillips (2003). “Neurochemical correlates of relapse to damphetamine self-administration by rats induced by stimulation of the ventral subiculum.” Psychopharmacology (Berl) 168(1-2): 99–108.

99. Taylor, J. R., P. Olausson, J. J. Quinn and M. M. Torregrossa (2009). “Targeting extinction and reconsolidation mechanisms to combat the impact of drug cues on addiction.” Neuropharmacology 56 Suppl 1(Suppl 1): 186–195.

100. Thibeault, K. C., M. Z. Leonard, V. Kondev, S. D. Emerson, R. Bethi, A. J. Lopez, J. P. Sens, B. P. Nabit, H. B. Elam, D. G. Winder, S. Patel, D. D. Kiraly, B. A. Grueter and E. S. Calipari (2025). “A Cocaine-Activated Ensemble Exerts Increased Control Over Behavior While Decreasing in Size.” Biol Psychiatry 97(6): 590–601.

101. Tierney, P. L., E. Degenetais, A. M. Thierry, J. Glowinski and Y. Gioanni (2004). “Influence of the hippocampus on interneurons of the rat prefrontal cortex.” Eur J Neurosci 20(2): 514–524.

102. Van den Oever, M. C., D. C. Rotaru, J. A. Heinsbroek, Y. Gouwenberg, K. Deisseroth, G. D. Stuber, H. D. Mansvelder and A. B. Smit (2013). “Ventromedial prefrontal cortex pyramidal cells have a temporal dynamic role in recall and extinction of cocaine-associated memory.” J Neurosci 33(46): 18225–18233.

103. Vertes, R. P. (2006). “Interactions among the medial prefrontal cortex, hippocampus and midline thalamus in emotional and cognitive processing in the rat.” Neuroscience 142(1): 1–20.

104. Vorel, S. R., X. Liu, R. J. Hayes, J. A. Spector and E. L. Gardner (2001). “Relapse to cocaine-seeking after hippocampal theta burst stimulation.” Science 292(5519): 1175–1178.

105. Wang, N., F. Ge, C. Cui, Y. Li, X. Sun, L. Sun, X. Wang, S. Liu, H. Zhang, Y. Liu, M. Jia and M. Yang (2018). “Role of Glutamatergic Projections from the Ventral CA1 to Infralimbic Cortex in Context-Induced Reinstatement of Heroin Seeking.” Neuropsychopharmacology 43(6): 1373-1384.

106. Wang, Q., J. Jin and S. Maren (2016). “Renewal of extinguished fear activates ventral hippocampal neurons projecting to the prelimbic and infralimbic cortices in rats.” Neurobiol Learn Mem 134 **Pt A**(Pt A): 38–43.

107. Warren, B. L., L. Kane, M. Venniro, P. Selvam, R. Quintana-Feliciano, M. P. Mendoza, R. Madangopal, L. Komer, L. R. Whitaker, F. J. Rubio, J. M. Bossert, D. Caprioli, Y. Shaham and B. T. Hope (2019). “Separate vmPFC Ensembles Control Cocaine Self-Administration Versus Extinction in Rats.” J Neurosci 39(37): 7394–7407.

108. Xu, H., L. Liu, Y. Tian, J. Wang, J. Li, J. Zheng, H. Zhao, M. He, T. L. Xu, S. Duan and H. Xu (2019). “A Disinhibitory Microcircuit Mediates Conditioned Social Fear in the Prefrontal Cortex.” Neuron 102(3): 668–682 e665.

109. Yang, S. S., N. R. Mack, Y. Shu and W. J. Gao (2021). “Prefrontal GABAergic Interneurons Gate Long-Range Afferents to Regulate Prefrontal Cortex-Associated Complex Behaviors.” Front Neural Circuits 15: 716408.

110. Zinsmaier, A. K., Y. Dong and Y. H. Huang (2022). “Cocaine-induced projection-specific and cell type-specific adaptations in the nucleus accumbens.” Mol Psychiatry 27(1): 669–686.

111. Zuo, Y., D. C. Smith and R. A. Jensen (2007). “Vagus nerve stimulation potentiates hippocampal LTP in freely-moving rats.” Physiol Behav 90(4): 583–589.

